# Correlated disasters and need-based transfers: The limits of risk pooling systems in simulated ecologies

**DOI:** 10.1101/230607

**Authors:** Marco Campenni, Lee Cronk, Athena Aktipis

## Abstract

Throughout their evolutionary history, humans have faced risks including drought, disease, natural disasters and other unexpected negative events. To deal with these risks, humans use a variety of risk management strategies, some of which involve relying on others in times of need in order to pool risk. However, the effectiveness of risk pooling strategies can be limited when there is high synchronicity of need. Here we investigate the limits of two resource transfer systems for pooling risk (need-based transfers, NBT, and debt-based transfers, DBT) in simulated ecologies with different degrees of correlated disasters using an agent-based model of the need-based transfer system of the Maasai. Overall, we find that survival is higher when shocks are less correlated among partners, when groups are larger, and when network structure is characterized by preferential attachment networks, which have a more modular structure than regular or small world networks. We also find that NBT strategies consistently outperform DBT strategies across a wide variety of parameter values and that the advantage of NBT over DBT is greatest when shocks are less correlated and group size is small. Our results also suggest that systems of sharing that are based on recipient need are less vulnerable than systems that are based on debt and credit, especially in small world and regular networks.

## INTRODUCTION

Risk is an unavoidable part of life for any organism living in a complex and changing environment. Humans have faced risks including drought, disease, natural disasters and other negative events throughout their evolutionary history. Part of the human solution to managing these risks is to help each other in times of need. Managing risk through being social is an ancient strategy - in fact the need to manage risk more effectively is one of the reasons that cells evolved to be multicellular, where they could share resources and engage in other social risk management practices (1). Thus, risk management is an adaptive problem that we have had to solve, not just since our human beginnings, but going back all the way to the very origins of life.

The risks that humans encounter can take many forms. Although some of these risks have positive expected values (e.g., windfalls to big game hunters in small scale societies; 2), many of them are negative. Every organism lives in an ecology that has some variation in resources over time and space, and many organisms live in environments where they encounter acute or chronic negative events. In human behavioral ecology the term risk refers to “unpredictable variation in an outcome with consequences that matter” (3).

Humans use several different strategies to manage risk (4). These include risk retention (i.e., accepting risk and absorbing losses), risk avoidance (i.e., reducing dependence on high variability outcomes), risk reduction (i.e., lowering the probability of or the size of losses), and risk transfer (i.e., moving risk from one party to another). In this framework, risk transfer is of particular interest for our understanding of human social behavior because it is the only risk management strategy that requires cooperation (2). One common form of risk transfer is risk-pooling (also referred to as risk-sharing). Risk pooling involves individuals sharing some of the risk of unexpected negative events. For example, among hunter-gatherers, if one individual has no food and another has more than they need as a result of successful foraging, the lucky individual may share with the unlucky individual, buffering that unlucky individual from the risks of unsuccessful foraging.

Humans are unusual as a species because they form long-standing, non-reproductive relationships with unrelated individuals, i.e friendships, and cooperation is a defining feature of these friendships (5). Humans also learn from and influence each other, showing a clear predisposition to cultural transmission (6). These facts contribute to the propensity of humans to form social networks, which can range in size from dozens to millions of people (7). Social networks show impressive structural regularities (7; 8), and both theoretical models and empirical results suggest that networks may have facilitated the development of large-scale cooperation in human beings (9; 10; 11; 12; 13).

To discover the possibly adaptive origins of human social networks and their relationship to cooperation, Apicella and colleagues (14) examined network features among Hadza foragers, whose way of life is thought to resemble that of our early ancestors (15). They find that cooperators tend to be connected to cooperators at both the dyadic and network level, conditions necessary to sustain cooperation (16).

Different network topologies can lead to different dynamics and different levels of resilience of the networks to perturbations. For example, Ash and Newth (17) demonstrated that preferential attachment networks (i.e, networks with modular components) are highly resilient to external perturbations and internal cascading effects. Modular components can help to isolate failures or, as in our models, shocks, and to reduce the overall complexity of the system. In modular networks, the consequences of local shocks are often propagated and resolved locally, affecting only a small part of the network and protecting most of the network from the perturbation. Less modular (e.g., regular networks and small-world networks) are more susceptible to the effects of shocks and show a lower resilience to them. In the present paper we explore how network topology interacts with the sharing system used by individuals in the network. We ask how vulnerable these networks are to collapse and whether network modularity is protective for both need-based and debt-credit based systems.

### Risk pooling systems can be based on need or on debt-credit relationships

This model is inspired largely by the system of risk pooling through need-based transfers among Maasai and other Maa-speaking pastoralists in East Africa called *osotua*, the literal meaning of which is “umbilical cord.” Osotua partners rely on their partners for help in times of need. For example, if a herder does not have enough livestock to support his family, he may ask one or more of his osotua partners for enough to bring his family up to the level necessary for survival. Transfers between osotua partners create neither credit nor debt, and there is no expectation that there will be an overall balance in total number of cattle transferred between partners, even over the long run. Transfers between osotua partners occur only in response to requests, and such requests must arise from genuine need and must be limited to the amount actually needed. Osotua relationships are imbued with respect, restraint, and a sense of great responsibility: in other words, they are sacred bonds (18, 2). Though the details vary, similar need-based transfer systems exist in many societies, from fisher-horticulturalists in Fiji to hunter-gatherers in Tanzania (2). These and other societies are part of The Human Generosity Project (www.humangenerosity.org), a large-scale transdisciplinary effort to study the biological and cultural influences on human cooperation through a combination of fieldwork, human subjects experiments, and computational modeling.

As part of The Human Generosity Project, we have designed and implemented several agent-based models that allowed us to explore the viability of various cooperative strategies for individuals living in different ecologies. This effort began with an agent-based model involving simulated pairs of osotua partners (19). In that model, herd survival was defined in terms of the agents’ success at keeping their herd sizes above a critical threshold, which was derived from the literature on herd demography and household food needs among East African pastoralists (20, 21). Agents whose herds dropped below the critical threshold for two consecutive rounds were removed from the simulation. Next we explored the viability of need-based transfers in simple networks that varied in size, connectedness and heterogeneity. We found that larger, more connected and more heterogeneous networks had higher herd survival. We also found that a selective asking rule, in which agents asked for help from their wealthiest partners, outperformed a random asking rule, in which agents asked for help from a randomly chosen partner (22).

Need-based transfer systems such as osotua are not the only form of resource transfer that can help to pool risk. Debt-based systems, such as those involving lending and credit, can also act as a safety net for individuals falling on tough times or needing resources for other reasons. In fact, the Maasai also have a debt-based transfer system called *esile*, which translates simply as “debt” (23). Esile exists side by side with the need-based transfer osotua system. Esile and osotua follow very different rules. In esile, unlike in osotua, credit and repayment of debts are the essence of the relationship: there is expectation from both parties that the debt will be repaid through an *elaata*, which means to set free or untie a knot (24; see also 2). Aktipis et al. (25) compared the need-based osotua system and the debt-based esile system in a dyadic interaction model, finding that need-based transfers lead to more risk pooling and higher survival than debt-based transfers. Moreover, need-based systems outperformed debt-based systems as the size and frequency of disasters increased. In the present paper we build on this previous work by placing agents in social networks with different features and exploring the impact of the synchronicity of disasters on the viability of these need-based vs. debt-based transfer systems.

Because many other pastoralist societies also have sharing rules that can be considered equivalent to osotua and esile (26; 27, 28; 29; 30; 31;32), we follow Aktipis et al. (25) and Cronk et al. (2) in using a more general, less Maasai-specific set of terms, referring to osotua as “need-based transfers” and esile as “debt-based transfers.” We operationalize need-based transfers and debt based transfers algorithmically in resource transfer rules used by the agents in our model (see Model description below).

### The limits of risk pooling

In the current paper, we investigate the limits of risk pooling in networks with different topologies and in environments with high or low synchronicity of need. Sharing systems based on risk pooling can help to buffer individuals from risks in the environment (see 33 for a network-based modelling approach; and,34 for an agent-based model of the influence that resource availability has on cooperation in the context of hunter-gatherer societies). However, if many individuals are simultaneously in need, then there may not be enough helping capacity in the system for individuals to recover from a negative event. When negative events strike many individuals simultaneously, it can be much more difficult for individuals to find others who are able to help because of the synchronicity of need. Among Maasai and other pastoralists, such a situation can arise due to a regional drought or epizootic disease.

The effectiveness of risk pooling can also be limited by the size and structure of the network - the more individuals there are to call on, the more likely individuals will be to find others who can help. But the structure of the network may also lead to unexpected vulnerabilities if shocks can reverberate and weaken the capacity of the network to future shocks. For these reasons, we varied both networks size and structure to explore the limits of both NBT and DBT strategies for pooling risk.

## METHODS

### Model

We used NetLogo (35) to model a population of agents in networks with varying topologies (Fig. 1). In this model, each node/agent in the network represents a household/family of approximately six individuals and each link represents a connection to another family in the network. Each agent begins with a herd of 70. The initial stock of 70 grows or shrinks during each time step of the simulation at a rate normally distributed around a mean of 3.4% (20). The maximum cattle herd size allowed in the model is 600, which represents a realistic approximation of the maximum cattle herd size for an averaged-sized household. During each time step there is also a chance of a loss (e.g., through a drought or a disease spreading in the herd). Following estimates of a family’s caloric needs and productivity in the dry season (20), we set the minimum size of a viable herd at 64 units as in (19, 22, and 25) (Fig. 2). Finally, across models we vary the probability, *p_shocks_*, that a loss is correlated among agents (i.e., it affects all agents at the same time) from 0 to 1 with steps of 0.1.

**Figure 1.**
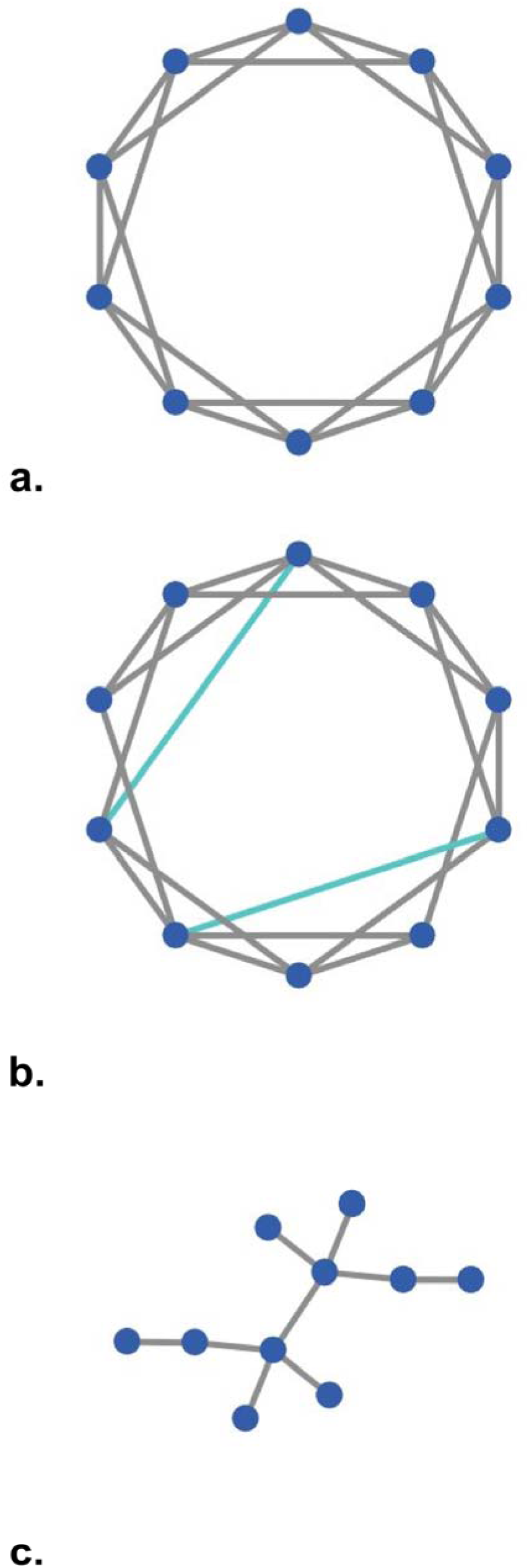
Examples of the network topologies used in our model. a) Regular network, b) Small-world network, and, c) Preferential attachment network.

**Figure 2.**
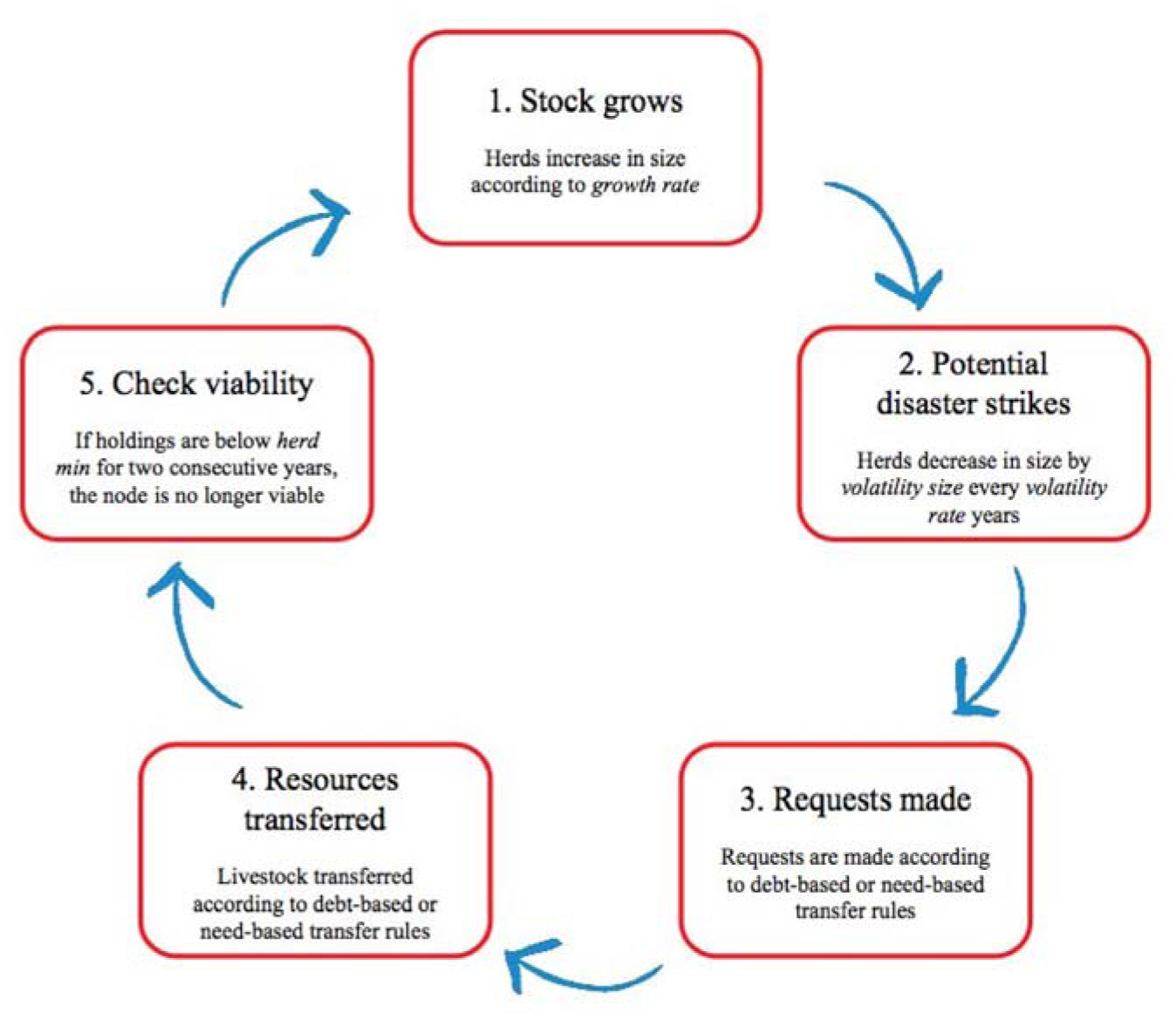
Model schedule. Each cycle of the model consists of five different steps or phases. In the first phase stock grows and each herd increases in size following the growth rate parameter. In the second phase potential disaster strikes may occur following the parameter volatility rate, and herds may consequently decrease in size following the parameter volatility size. In the third phase a request is made by agents in need according to the strategy they adopt. In the fourth phase resources may be transferred according to the strategy adopted by agents providing help. Finally, in the fifth phase, a viability check is performed to remove from the model agents with herd size below the sustainability threshold of 64 for two consecutive rounds (or years).

We simulated both need-based and debt-based livestock transfers between individuals by implementing these rules as algorithms that could be employed by each agent.

### Need-based transfers

If you drop below the critical threshold for livestock holdings, ask your wealthiest partner for help (i.e., agents used a selective asking rule rather than a random asking rule as in 22); if you are asked for help and can afford to provide it without putting yourself below the critical threshold for survival, do so.

### Debt-based transfers

Debt-based transfer agents also asked for help when they were in need. But otherwise they differed from need-based transfer agents. Debt-based agents only transfer cattle if they are asked by a partner who is in good standing. These debt-based agents kept track of the amounts they owed to and were owed by the other agents in their networks. Recipients of loans repaid these loans as soon as they had enough livestock to do so without going below the sustainability threshold of 64 units. If five rounds went by after a transfer without repayment, then the agent who gave the loan considered this partner to no longer be in good standing and therefore would not provide a loan to this agent in the future.

In the appendix we provide an algorithmic description of both transfer strategies.

##### GLOSSARY

###### Regular network

In graph theory, a regular graph (or network) is a graph where each node has the same number of neighbors. In other words, every node has the same degree (i.e., number of connections with other nodes).

##### Small-world network

A small-world network is a network where the typical distance, L, between two randomly chosen nodes (the number of steps required) grows proportionally to the logarithm of the number of nodes N in the network (Watts & Strogatz 1998). In the context of a social network, this results in the small world phenomenon of strangers being linked by a short chain of acquaintances. Many empirical graphs show the small-world effect, e.g., social networks, the underlying architecture of the Internet, wikis such as Wikipedia, and gene networks.

##### Preferential attachment network

Preferential attachment means that the more connected a node is, the more likely it is to be connected to other nodes. Intuitively, preferential attachment can be understood if we think in terms of social networks connecting people. Here a link from A to B means that person A “knows” a person B. Heavily linked nodes represent well-known people with lots of relations. When newcomers enter a community, they are more likely to become acquainted with one of those more visible people rather than with a relative unknown. The Barabási-Albert (BA) model is an algorithm for generating random scale-free networks using a preferential attachment mechanism. Several natural and human-made systems, including the Internet, the world wide web, citation networks, and some social networks are thought to be approximately scale-free and certainly contain few nodes (called hubs) with unusually high degree as compared to the other nodes of the network.

##### Degree

The degree of a node in a network is the number of connections or edges the node has to other nodes.

##### Modularity

Modularity is one measure of the structure of networks or graphs. It was designed to measure the strength of division of a network into modules (also called groups, clusters or communities). Networks with high modularity have dense connections between the nodes within modules but sparse connections between nodes in different modules. Preferential attachments networks have higher modularity than random networks and small world networks.

## RESULTS

We investigated the limits of two resource transfer systems for pooling risk (need-based transfers, NBT, and debt-based transfers, DBT) in simulated ecologies with different degrees of correlated disasters using an agent-based model of the resource transfer systems of the Maasai. First, we compared two different strategies, need-based transfers, NBT, and debt-based transfers, DBT, in environments with different likelihoods, p_shocks_, of correlated shocks. Next, we varied group sizes from six to one hundred, reflecting plausible values for actual group sizes in actual small scale societies.

Finally, we varied network topologies. The network topology is the specific social structure governing the relationships in a group of individuals and affecting their behaviors. We investigated sharing behaviors in three main network topologies: a regular network, a small-world network, and a preferential attachment network (for details about the algorithmic implementation of the different topologies and their meanings, see the Methods section). Throughout, we measured survival at 100 rounds, corresponding to 100 simulated years.

### NBTs outperform DBTs when the correlation of shocks is low

We investigated the effects of synchronicity of shocks by varying the correlation of shocks, p_shocks_ from a scenario where all shocks are uncorrelated (i.e., idiosyncratic scenario) to a scenario where all shocks are completely correlated (i.e., systemic scenario) and affect all agents at the same time. This probability, p_shocks_, represents the probability between 0 and 1 of a shock to affect all members of a group (in our model of a social network of agents) at the same time. Probability p_shocks_ = 0 models the situation where a shock is very likely to affect just a single agent in the network at time t. Probability p_shocks_ = 1 models the scenario where all agents from the same network are affected by the shock at time t (see model description in Method section for details). We also considered intermediate scenarios varying the probability of correlated shocks from 0 to 1 by steps of 0.1. At each step, we noted the median survival of agents over 10,000 replicates of the model. Results from our models show (see Fig. 3) that increasing the probability p_shocks_ of correlated shocks progressively decreases the survival of the system and the gap between adopting NBT strategy and DBT strategy (p-values of Kruskal-Wallis Test < 0.05 for N = 6 and p_shocks_ < 0.6). Thus, the resilience of the network strongly depends on this parameter. Moreover when p_shocks_ = 0 and N = 6, the need-based transfer strategy is the only one that allows the system to avoid a collapse (compare results a and b shown in Fig. 6 when p_shocks_ = 0 and N = 6). When p_shocks_ is increased, networks adopting the need-based transfer strategy outperform those that adopt the debt-based transfer strategy. When p_shocks_ is greater than 0.5, the system quickly collapses (at round 50 or before) regardless of whether individuals use need-based on debt-based transfer strategies.

**Figure 3.**
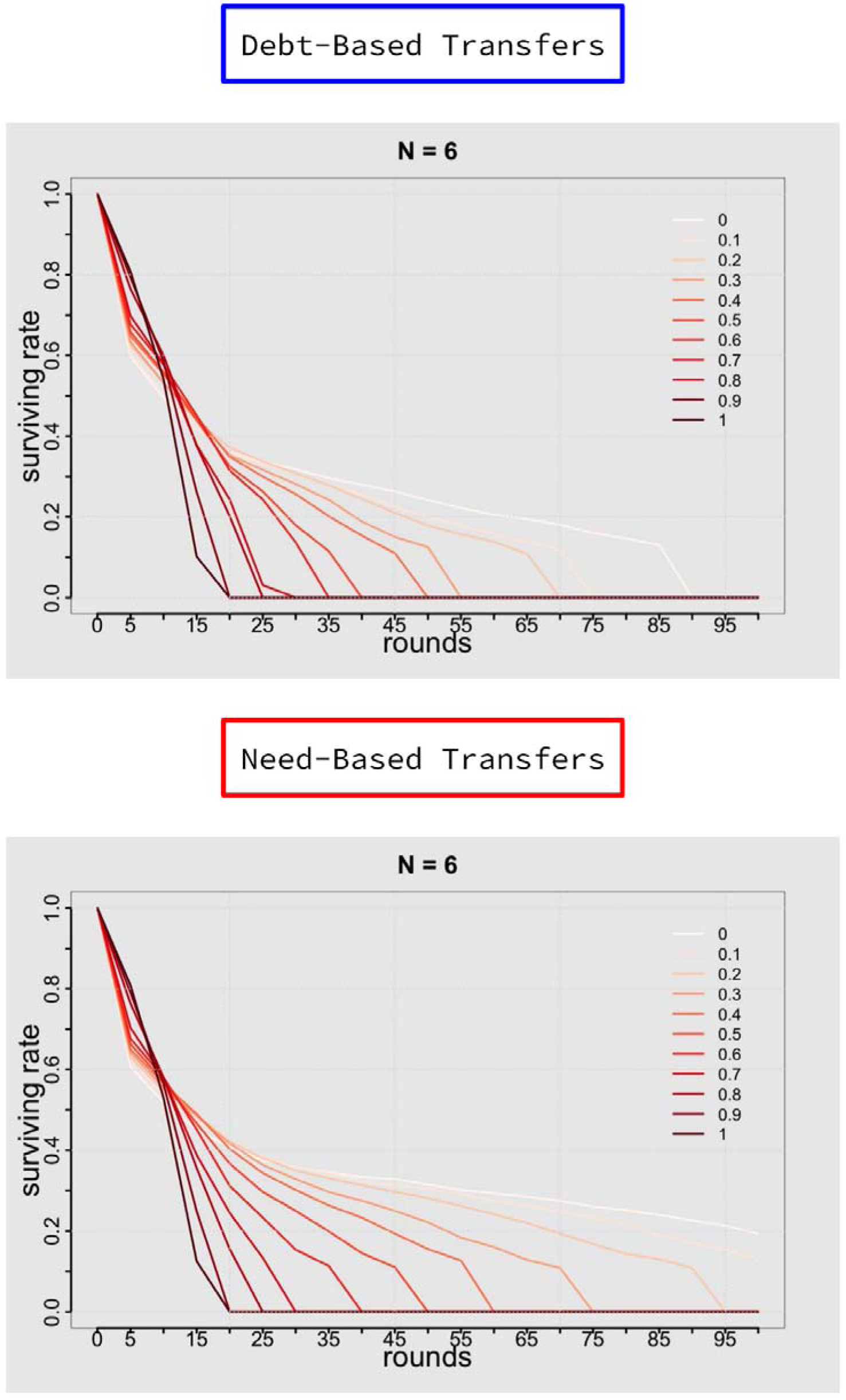
The figure shows the survival of the system (y-axis) as a function of the correlation of shocks. The darker the red line, the higher the correlation of shocks. NBT systems outperform DBT systems most dramatically when the correlation of shocks is low (lighter red line). When correlation of shocks is high (dark red lines), no systems survive to the end of the simulation. When correlation of shocks is low, some NBT systems are able to survive until 100 simulated years, while DBT systems do not.

**Figure 6.**
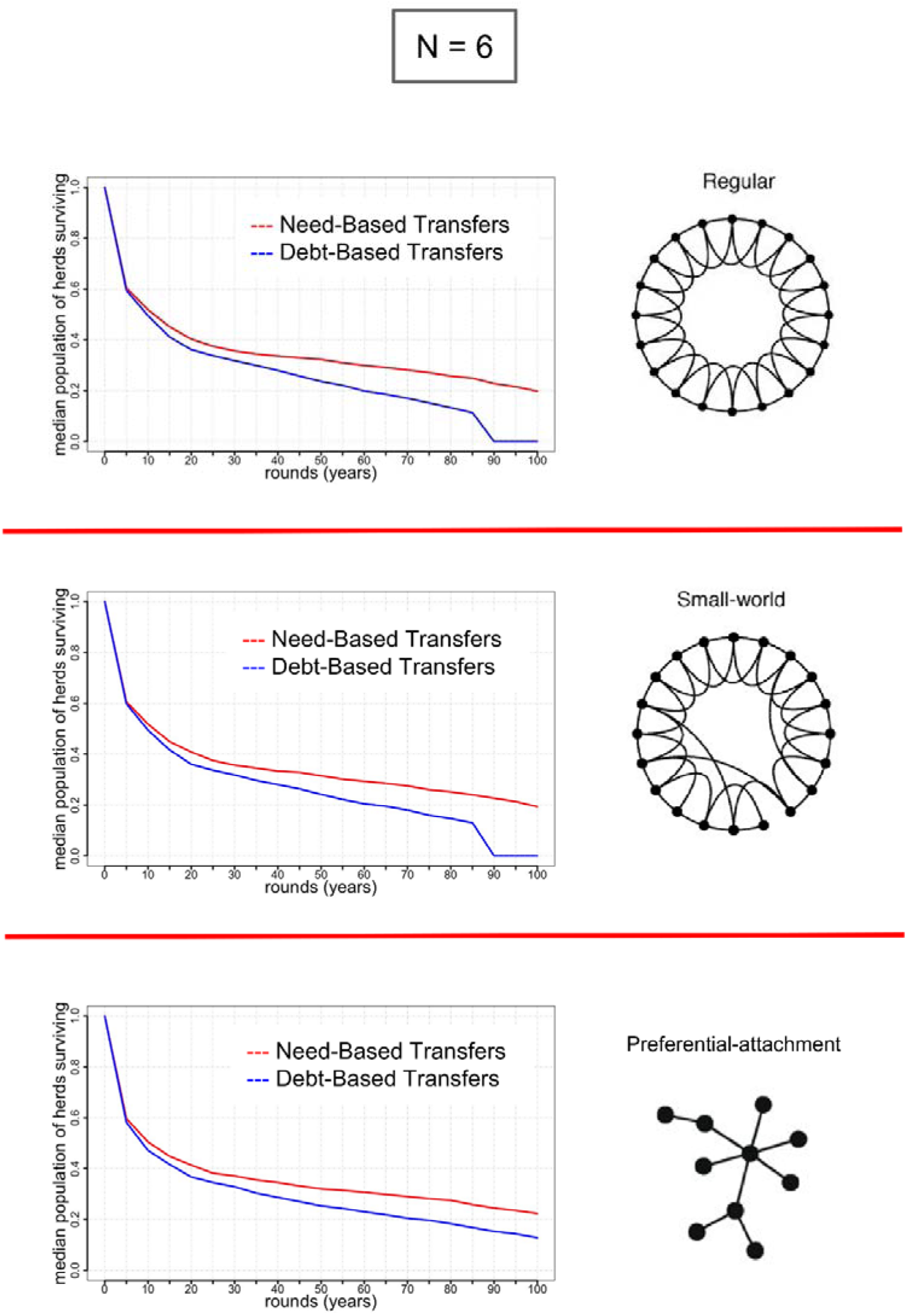
The survival of the system (y-axis) as a function of time (x-axis), considering three different network topologies and no correlation of shocks. (a) In regular networks, (b) in small-world networks, and, (c) in preferential attachment networks NBT strategies outperform DBT strategies. DBT strategies are most successful when networks have a more modular structure (i.e., preferential attachment networks, figure c), though NBT systems still outperform DBT systems in modular networks.

### When group size is small NBTs outperform DBTs

We kept constant p_shocks_ and varied group size N to see how the network size affects the ability of the system to avoid collapse. Small networks (N=6) of agents adopting the DBT strategy collapse even when there is a low correlation of shocks (i.e., p_shocks_ = 0.1), while NBT networks survive shocks at this correlation level. When N=10, NBT systems exhibit better performances in absorbing shocks (p-values of Kruskal-Wallis Test < 0.05), but systems adopting the DBT strategy can survive, too. Finally, although starting from N=10 DBT strategy networks can survive shocks (see Fig. 4), statistical analysis shows that there is a difference in performances of networks adopting NBT strategy and those adopting DBT strategy (Kruskal-Wallis Test shows that p-values < 0.05 for N < 90). This means that, as the size N of the network increases, the DBT strategy becomes more viable and comparable to the NBT strategy, but NBT strategy networks perform better than DBT strategy networks when the group size is relatively small (i.e, N < 90).

**Figure 4.**
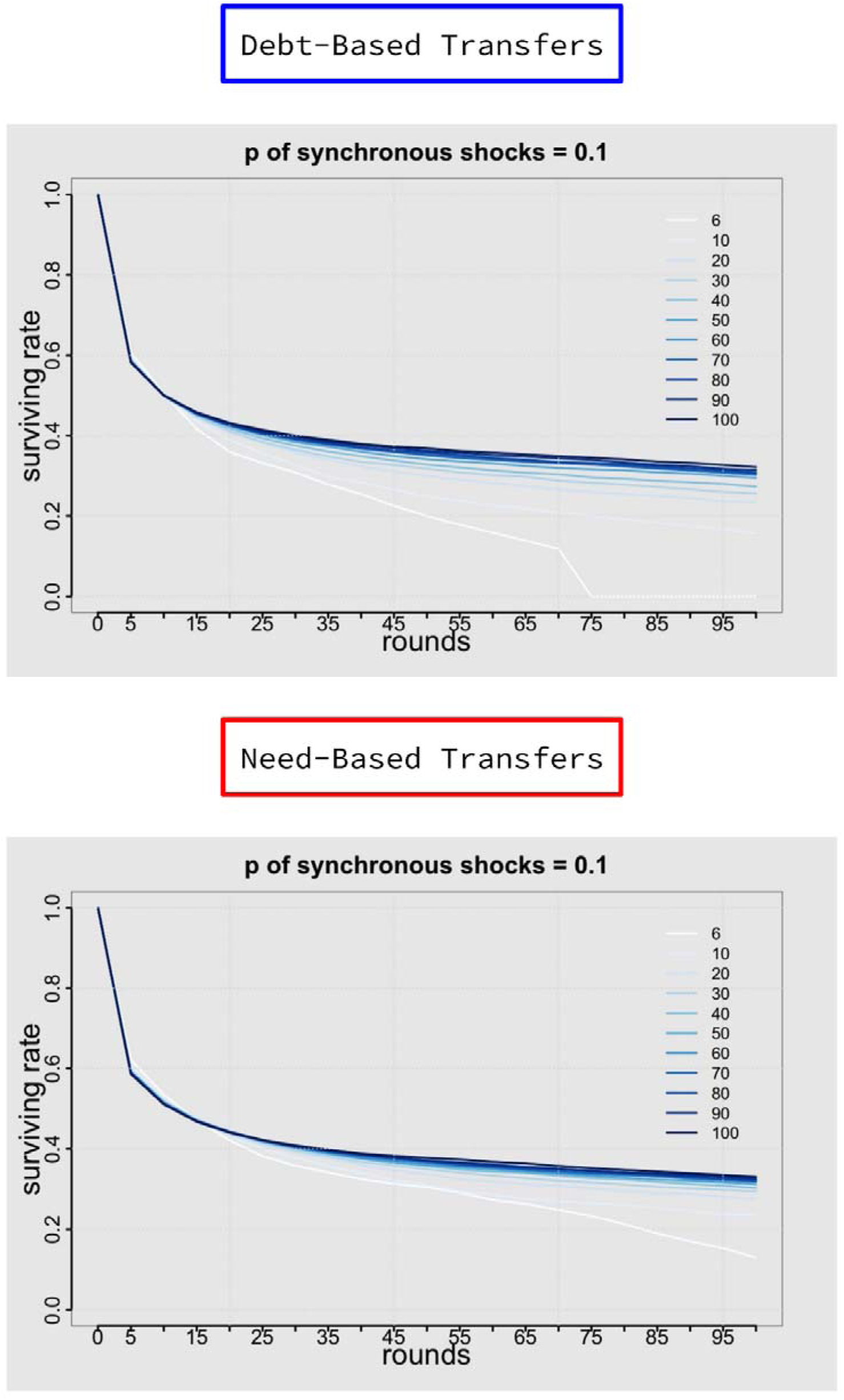
The figure shows the survival rates for individuals in the system (y-axis) as function of the size of the group. The darker the blue line, the bigger the group. NBT systems outperform DBT systems when the group is small. When group size is small, some NBT systems can survive past 100 simulated years, but DBT systems cannot.

### When group size is small and correlations of shocks is low NBT has the greatest advantage over DBT

We compared the survival of need-based transfer systems and debt-based transfer systems while covarying both the probability, p_shocks_, of correlated shocks and the group size N to quantify the resilience of both strategies to shocks. Fig. 5 shows that small networks (N=6) that adopt the NBT strategy exhibit greater resilience than systems that adopt the DBT strategy. More specifically, when p_shocks_ < 0.2, NBT systems survive, while DBT systems do not. When p_shocks_ ≥ 0.2, both systems collapse before the end of the simulation. Increasing the size of the network reduces the gap between NBT system and DBT system performances. For N = 30, NBT systems outperform (Kruskal-Wallis Test p-values < 0.05) DBT systems when p_shocks_ < 0.5; when p_shocks_ = 0.6 both systems collapse. For N = 50, NBT systems performs better than DBT systems if and only if p_shocks_ < 0.4; for values of p_shocks_ between 0.4 and 0.7 it performs equally well. Finally, for p_shocks_ > 0.7 both systems collapse. For N ≥ 50, the advantage in adopting the NBT strategy is limited to very low p_shocks_ (i.e., p_shocks_ ≤ 0.2), but generally, any system adopting either NBT or DBT strategy is much more resilient to correlated shocks: in fact, in both cases, NBT and DBT systems avoid the collapse for pshocks values ≤ 0.7.

**Figure 5.**
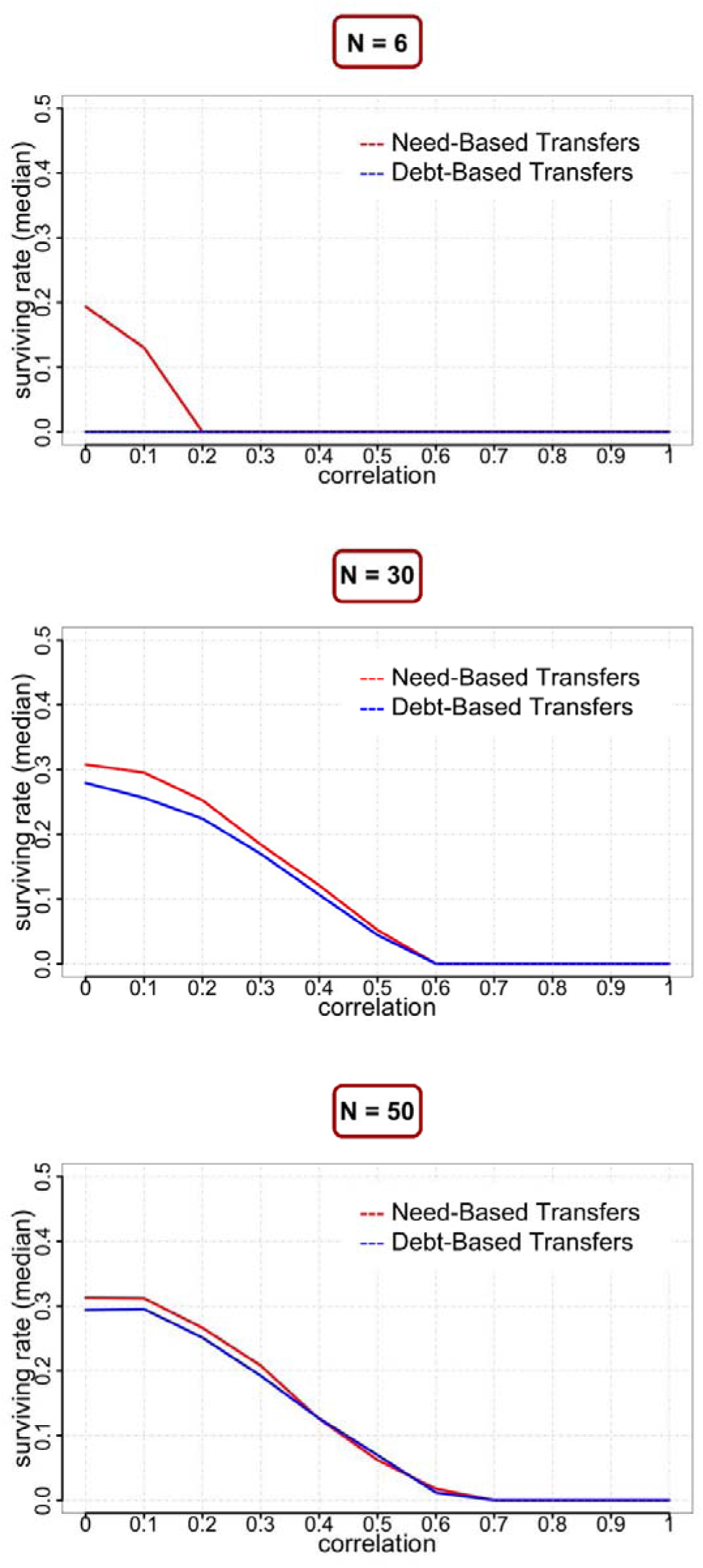
The figure shows how the survival of the system (y-axis) adopting either NBT or DBT may vary as function of the correlation of shocks (x-axis), varying the group size of the network, considering N=6 (a), N=30 (b), and, N=50 (c). NBT systems outperform DBT systems when the network is small and the correlation of shocks is low (e.g., left side of figure a).

### NBTs outperform DBTs across all network structures and modular networks are most resilient

In the baseline scenario where there is no correlation of shocks, the network size includes only six nodes, and the network topology is either regular or small-world, the need-based transfer strategy performs better than the debt-based transfer strategy (see results **a** and **b** in Fig. 6). In both cases, networks of agents adopting debt-based rules collapse after 90 rounds. In the case of preferential-attachment networks, however, both strategies allow the system to survive, and the difference between the performance of need-based transfer systems and debt-based transfer systems is smaller than for other networks (see result **c** in Fig. 6).

Both NBT systems and DBT systems show higher survival across all levels of correlated shocks when the network is modular. Modules in (complex) networks are sub-groups of nodes that are highly interconnected within, and loosely connected outside the group. DBT systems do collapse (no one survives after 85 simulated years) in normal and small networks, but they do not collapse in preferential-attachment networks (at least, not within the time window we considered, i.e., 100 simulated years).

## Discussion

Humans have faced risks including drought, disease, natural disasters and other unexpected negative events throughout their evolutionary history. To deal with these risks, humans use a variety of risk management strategies, some of which involve relying on others in times of need in order to pool risk. However, the effectiveness of risk pooling strategies can be limited when there is high synchronicity of need. Overall, we find that survival is higher when the correlation of unexpected events is low, which enables those unaffected by the shock to help those in need. Moreover, the survival is higher when groups are larger because there are more individuals able to share resources and absorb the negative effects of unexpected events. Finally, the structure of social network matters: the more modular the structure is, the more resilient it is to external perturbations.

The superiority of need-based transfers over debt-based transfers in these models can be attributed to the simple fact that debt-based agents sever ties with other agents who, particularly in a volatile environment, might actually be helpful to them at some point in the future. Need-based agents, in contrast, do not expect to be repaid and so maintain all their partnerships, thus increasing the likelihood that they will receive help when they need it. This leads to the question of why need-based and debt-based transfers coexist. We propose that each one is appropriate in specific but quite different circumstances. When needs arise on a regular, predictable basis, it is possible for people to agree to a balanced exchange of favors: You help me today, I’ll help you tomorrow. When such favors are not repaid, it also makes sense to end the relationship and try again with someone else. But when needs arise unpredictably, as in our models, it is sensible to simply help those in need so that they will still be around to help you if you yourself are in need at some point in the future. This contrast is well illustrated by a pattern observed among ranchers in the American Southwest (36, 2). For help with predictable things, such as rounding up livestock for branding or marketing, ranchers agree to trade favors in the form of skilled labor. But when needs arise unpredictably (e.g., injuries and equipment failures), they provide help to one another with no expectation of any repayment beyond a similar kindness should they themselves ever suffer from a similarly unpredictable need.

Another advantage that need-based transfer systems may have over debt-based systems is the sheer simplicity of need-based transfer rules: Ask if you need, give if you can. The rules underlying a debt-based system, in contrast, are far more complex, involving not only rules about debt, credit, and repayment but also the maintenance of memories regarding the current status of each relationship one has. Need-based transfer agents, in contrast, simply need to keep track of whether their own resource holdings are above their survival threshold and occasionally calculate whether they can afford to help a partner in need. The low cognitive requirements of need-based transfer systems suggest that they may have predated account-keeping in our species’ evolutionary history and could be more phylogenetically widespread than systems requiring the tracking of credits and debts (25).

One of our main findings is that correlated disasters lead to much lower survivorships, regardless of which rule of property transfer agents may use, than uncorrelated disasters. This finding is similar to one based on a network model of shocks in financial systems (37). In that simulation, the resilience of banking networks to systemic shocks is very low compared to their resilience to idiosyncratic shocks. The authors argue that the detrimental impact of systemic shocks comes from the fact that the shocks themselves reduce the wealth of each node of the network system at the same time, which makes them more vulnerable to shocks in subsequent periods. Hence, large shocks are likely to be followed by cascading defaults, thereby destabilizing the whole system.

One way to reduce the correlation of shocks is to scale up the system. If shocks happen in specific locations, then pooling risk more broadly can reduce the synchronicity of shocks within the system over all. In societies with modern infrastructure for transportation and communication and with institutions (e.g., states, corporations, and non-profits) that routinely operate on large scales, such scaling up may be relatively easy. In societies with subsistence economies, scaling up may be more difficult. However, it may not be entirely impossible. For example, Fijian islanders adjust the social distance between themselves and the person they ask for help depending on whether the need is very local or more widespread. For example, when illnesses and injuries occur, people ask those close at hand, such as close kin, but when widespread disasters such as droughts and cyclones strike, they draw upon ties to distant kin in other villages where the impact of the disaster may be less severe (2).

Whenever people cooperate at some cost to themselves, there is the potential for cheating as people seek to reduce the costs they pay or increase the benefits they receive. In a need-based transfer system, cheating consists of feigning either need or an inability to help those in need. Among Maasai and other pastoralists, such cheating is made difficult by the fact that the primary form that wealth takes is publicly visible: Livestock. In addition, osotua relationships are imbued with a sense of sacredness that makes cheating virtually unthinkable (18). Nevertheless, it does behoove people to choose osotua partners very carefully. Partner choice is one possible way to enhance the assortment of cooperators with one another, and it can be implemented through a variety of behavioral rules for choosing and maintaining relationships (38, 39, 19; 40; 41; 42; 43). Among the Maasai, the selection of osotua partners resembles a courtship process, with prospective osotua partners getting to know each other and giving small gifts over a period of years. When a degree of trust has been established, the relationship may then be recognized as osotua. In future models of need-based transfers on networks, we plan to model this relationship formation process explicitly to investigate whether partner choice increases the viability of the need-based transfer strategy.

Across many systems, modularity has been shown to increase resilience to perturbations in networks. This is the case for the evolution and stability of biological networks (17, 44), the scalability and efficiency of large-scale infrastructure (45, 46), and the development of economic and social systems (47, 48). In addition, one of the most striking characteristics of the networks resulting from an optimization process aimed at maximizing resilience against cascading failures is their modular design (17). Self-contained or modular components are often used as a means to isolate failures or, as in our models, shocks, and reduce the overall complexity of the system. Modularity in complex networks could also be serving a similar function by limiting the effects of shocks on the system as a whole. Networks showing a lower degree of modularity (e.g., regular networks and small-world networks) are more vulnerable to the effects of shocks.

We found that the most modular network structure, the preferential attachment network, was associated with the highest survival for both need-based and debt-based transfer strategies. The preferential network structure also allowed the debt-based strategy to survive in conditions that were otherwise impossible (when correlation of shocks is high). Future work can investigate in greater detail which characteristics of preferential attachment networks are driving this effect. Our preliminary analyses show that preferential attachment networks have higher degree (i.e, the number of connections or edges the node has to other nodes), higher Eigenvector centrality (i.e, a measure of the influence of a node in a network) and, of course, higher modularity (i.e, the division of a network into modules, groups, clusters or communities). In future work, the relative importance of each of these features could be investigated in greater depth.

## Conclusion

Our findings may have implications for real-world disaster scenarios. Not surprisingly, when disasters strike entire networks at the same or nearly the same time, networks rarely survive. However, when disasters occur asynchronously, survival is improved if agents use a need-based transfer strategy rather than a debt-based strategy, if networks are larger, and if networks are more modular. Because it may be difficult in many circumstances to simply create larger or more modular networks, perhaps the most important take-home message is that, when disasters and other negative events arise unpredictably, it is more adaptive to simply give to those in one’s network who are need if one is able to do so, and to do so without any expectation of repayment, rather than to give with the understanding that the relationship will end if the gift is not repaid. Spontaneous helping networks often emerge in the aftermath of disasters, with individuals helping strangers without expecting anything in return (49; 50). For example, after the 1906 earthquake in San Francisco, even traditionally market driven exchanges, such as food deliveries and transportation became need-based systems, with food being distributed and public transportation running for free until after the crisis passed (50). These examples, combined with the modelling results we report above, suggest that need-based helping systems may an important part of the human toolkit for surviving during correlated disasters.

The authors declare no competing financial interests.

For additional analyses and details, see Appendix.

## Authors’ contributions

MC, LC, and AA conceived of the study, designed the study, coordinated the study and helped draft the manuscript. MC created the original model, accomplished the simulations and carried out the statistical analyses. All authors gave final approval for publication.

## Acknowledgments

This material is based upon work supported by the National Science Foundation under Grant No. SES-0345945 to Arizona State University’s Decision Center for a Desert City (DCDC) and Grant No. BCS-1324333 to Cronk and Iyer, National Institute of Health Grant No. F32 CA144331, and a grant from the John Templeton Foundation. We thank the Institute for Advanced Study in Princeton, the Center for Theological Inquiry in Princeton, the Center for Advanced Study in the Behavioral Sciences at Stanford University and the Wissenschaftskolleg in Berlin. We would also like to thank participants in the National Evolutionary Synthesis Center catalysis meeting for Synthesizing the Evolutionary and Social Science Approaches to Human Cooperation, the members of the Human Generosity Project and the members of the Cronk and Aktipis lab groups. Any opinions, findings, conclusions, or recommendations expressed in this material are those of the authors and do not necessarily reflect the views of the National Science Foundation (NSF), the National Institute of Health (NIH), or the John Templeton Foundation.

## Funding

This material is based upon work supported by the National Science Foundation under Grant No. SES-0345945 to Arizona State University’s Decision Center for a Desert City (DCDC) and Grant No. BCS-1324333 to Cronk, National Institute of Health Grant No. F32 CA144331, and a grant from the John Templeton Foundation to Aktipis and Cronk.

## Appendix.

Need-based transfer rules were implemented as follows (consistently with Aktipis et al. 2016, Aktipis et al. 2011, and Hao et al. 2015):

1. **Need-based asking rule**: Individuals ask their partners for livestock only if their current holdings are below the asking threshold (i.e., the minimum stock size of 64).
2. **Need-based giving rule**: Individuals give what is asked, but not so much as to put their herds below the giving threshold (also the minimum stock size of 64).

Debt-based transfer rules were implemented as follows:

1. **Debt-based payback rule**:
  a. If livestock have been previously transferred from the partner to the actor and the actor has enough to pay back without going below sustainability threshold (*resource min*), the actor ‘pays back’ livestock to his partner according to the actor’s *repayment probability*
2. **Debt-based partner credit check rule**:
  a. Checks whether partner is in good standing, which includes not having exceeded *tolerated delay* or *credit size* (when applicable)
3. **Debt-based asking rule**:
  a. As with the need-based transfer asking rule, individuals ask their partners for livestock if their current herd size is below the sustainability threshold of 64.
4. **Debt-based giving rule**:
  a. Response to partner. If a request is made, actors give if two conditions are met:
    i. If no debt remains from a previous request and partner is in good standing (meaning that previous debt had not existed for longer than *tolerated delay*)
    ii. The amount transferred cannot exceed the *credit size* extended to the partner

### Network Generation

To create a baseline for all comparisons, we decided to first implement a **homogeneous regular** network of agents to test the average herd survival of populations adopting either a need-based transfer strategy or a debt-based transfer strategy. By homogeneous we mean that all individuals in the network have the same number of partners. Each vertex in the network represents a family and each pair of partner families is connected by an edge. N represents the network size (i.e., the number of families in the network); k represents the average degree of the network, i.e., the average number of partners per family. Each relationship is bidirectional, thus each node i, (1 ≤ i ≤ N) has degree four (d_i_=4) and is connected to two of its closest neighbors from both sides.

We then generated **small-world** networks and **preferential attachment** networks to investigate the effects of network topologies on performances of both strategies.

The **small-world networks** were implemented starting from regular networks and randomly rewiring a small fraction (β = 0.1) of the edges of a homogeneous network. A **small-world network** is a type of network in which most nodes are not neighbors of one another, but the neighbors of any given node are likely to be neighbors of each other; moreover, most nodes can be reached from every other node by a **small** number of steps.

The **preferential attachment** networks were generated adapting to our purposes the code from Wilensky (2005). The network generation algorithm starts with two nodes connected by an edge. At each step of the algorithm, a new node is added. Each new node picks an existing node to connect to randomly, but the node’s chance of being selected being directly proportional to the number of connections it already has (i.e., the number of edges linking it to other nodes), or its "degree." This is the mechanism which is called "preferential attachment."

The networks that result from running this algorithm are often called "scale-free" or "power law" networks. Those networks do not present a normal distribution of the number of connections of each node - instead they follow a power law distribution. Power law distributions are different from normal distributions in that they don’t have a peak at the average, and they are more likely to include extreme values (see Albert & Barabási 2002 for a further description of the frequency and significance of scale-free networks).

Need-based transfer asking rule: agents (i.e., nodes in the network) make a request for cattle only if their current holdings are below the critical threshold (i.e., the minimum herd size of 64). Each agent is allowed to make only one request per year (or round). Need-based transfer giving rule: agents (i.e., nodes in the network) give what is asked, but not so much to make their cattle holdings reach the critical threshold (i.e., the minimum herd size of 64); if the request R is higher than the amount A necessary to reach the critical threshold, the agent will give just A.

Debt-based transfer asking rule: agents (i.e., nodes in the network) make a request for cattle only if their current holdings are below the critical threshold (i.e., the minimum herd size of 64). Each agent is allowed to make only one request per year (or round).

Debt-based transfer giving rule: agents (i.e., nodes in the network) give what is asked only if the partner agent asking for help is a trustworthy partner (i.e., the partner already paid any debit), but not so much to make their cattle holdings reach the critical threshold (i.e., the minimum herd size of 64); if the request R is higher than the amount A necessary to reach the critical threshold, the agent will give just A.

Asking order: agents are randomly selected to make request each year (or round).

Asking strategy: we implemented a “selective” asking strategy in a way that, assuming that all agents have complete information about the herd size of their partners, agents are able to select the wealthiest partner among all their osotua or esile partners to make a request. When two or more of an agent’s partners have the same maximum herd size, the selection will be randomly performed with equal probability.

### Simulation

Each simulation (or run of the model) consists of five phases (see Figure 2 in MS for a graphical representation and detailed description of phases): Initialization, Random growth, Disaster, Request, Response, and Viability check. After initialization, during each phase agents are randomly selected to act, interact or simply experience changes in their livestock. We varied the size of the network considering networks of N={6,10,20,30,40,50,60,70,80,90,100} nodes. For homogeneous regular networks the average degree is k=4. Finally, in the simulations, the herd size H_i_^(n)^ of each agent i (1 ≤ i ≤ N) at year t, is updated at the end of year t simultaneously and is tracked until t = 100. To compensate for the stochasticity of random initial conditions, for every network topology (i.e, regular, small-world and preferential attachment networks), simulations were replicated 10000 times and the average values are reported.

We then compared the performance over time of populations of need-based transfer agents with other types of populations of debt-based transfer agents. Additional details regarding the model schedule, parameter values and model design can be found in the ODD protocol description of the model that follows.

### Initialization

Each network topology is specified by an N x N adjacency matrix *A* = {*a*_*ij*_}. Because the reciprocal nature of need-based transfer and debt-based transfer relationships as we implemented in the model, the network is undirected, i.e., if a node i is a partner node to another node j, then *a_ij_* = *a_ji_* = 1. Consequently, the adjacency matrix A is a symmetric matrix with row sum k.

For a **homogeneous regular** network, A has permutation symmetry (i.e., each node is identical) and additional symmetries depending on the structure of the edges of the network.

A **small-world** (in our simulations β = 0.1) Watts-Strogatz network with a given *β* is generated randomly selecting 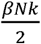 edges, disconnecting one end and reconnecting the link to another different randomly selected node. 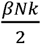 pairs of subscripts (i,j) are randomly selected from {(i,j)|A(i,j)} with equal probability. The new adjacency matrix *Â* will lose the permutation symmetry but the row sums will remain unchanged.

A **preferential attachment** network generation begins with an initial connected network of m_0_ nodes. Then, new nodes are added to the network one at a time. Each new node is connected to m ≤ m_0_ existing nodes with a probability p that is proportional to the number of links that the existing nodes already have. Formally, the probability p, that the new node is connected to node i is 

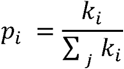

where k_i_ is the degree of node i and the sum is made over all pre-existing nodes j (i.e. the denominator results in twice the current number of edges in the network).

Strongly linked nodes (i.e., so called "hubs") tend to quickly accumulate even more links, while nodes with only a few links are unlikely to be selected for creating a new link. The new nodes have a "preference" to attach themselves to the already well connected nodes.

For all network topologies, initial herd size are set to H_1_^(0)^=70, 1 ≤ i ≤ N, for all agents at the beginning of each simulation.

### Random growth

We implemented the same growing dynamics implemented in Aktipis et al. 2011, Hao et al. 2015 and Aktipis et al. 2016. Thus, each year an agent’s herd i grows at random rate g_i_^(n)^ which is sampled from a Gaussian distribution. The growth rate distribution is given by

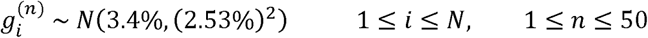

meaning that the growth rate follows a Gaussian distribution with mean 3.4% and a standard deviation 2.53%, which represents a typical annual growth rate for cattle herds in East Africa (Dahl and Hjort, 1976). The growing process implemented in the model implies that before considering shock events that may happen over the same year, at the end of year n the herd size for an agent i is

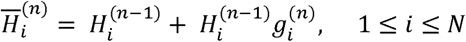

The Gaussian distribution of growth rate implemented as such gives us also a very small probability of negative growths, which means that independently of shocks (e.g., drought, disease, theft) in some years deaths may exceed births.

### Disasters

At each single cycle of the model, the probability of correlated shocks p_shocks_ determines whether the size of the shock affects a single agent or all agents at the same time. This is implemented by a simple comparison between the current value of p_shocks_ and a value v randomly selected between zero and one; if v < p_shocks_, then the shock will affect all agents at the same time, otherwise, the shock will affect only the current agent. Then, the actual happening of shocks is modeled by a Poisson process. We assume that the sequence of shocks is generated by a Poisson process and that the time until the next disaster happens is exponentially distributed with mean 0.1.

In the case of completely uncorrelated shocks (i.e., p_shocks_ = 0), each agent’s herd is considered independent and each year there is a probability p = 0.1 that a shock happens. If the current year n is a disaster year for agent i, thus a random number 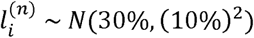 is drawn to quantify the percentage of the herd that is lost in this year.

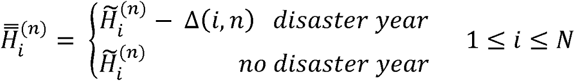

### Request

After accounting for random growth and disaster losses, the model selects for agents whose herd sizes fall below the critical threshold of sustainability (*θ* = 64); then, each of those agents are randomly selected for asking their wealthiest partner for help. Finally, a request for enough cattle to bring the herd size back to the threshold is performed

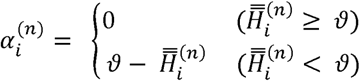

Where α_i_^(n)^ is the number of cattle asked by agent i.

Each agent is allowed to make only one request per year (or round), even if no livestock are received as a result of the request.

### Response

#### Osotua

Agents respond to osotua requests one a time following the order in which the requests are made.

When an agent j is asked by another agent i, j will respond by giving *γ_j→i_*^(*n*)^ cattle to i where

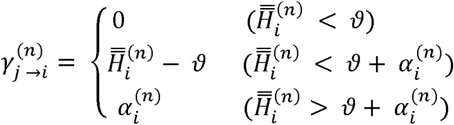

In other terms, if j has more cattle than θ, then it will fully respond to the request made by i. If j has more than θ + α, then it will fulfill the request made by i giving α_i_^(π)^ cattle to i; otherwise, j only gives all cattle exceeding the threshold θ to i.

#### Debt-based transfer

Agents respond to debt-based transfer requests one a time following the order in which the requests are made.

When an agent j is asked by another agent i, j will respond if and only if the agent i results to be a trustworthy partner (i.e., agent i already gave back to agent j the amount k received during past livestock transfers); j will respond by giving *γ_j→i_*^(*n*)^ cattle to i,
Where

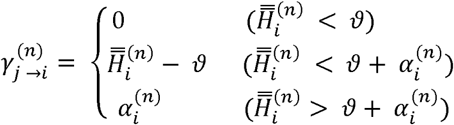

In other terms, if j has more cattle than θ, then it will fully respond to the request made by i. If j has more than θ + α, then it will fulfill the request made by i giving α_i_^(n)^ cattle to i; otherwise, j only gives all cattle exceeding the threshold θ to i.

### Viability check

When the “resources transferred” phase ends, the herd sizes are finalized for the current year,

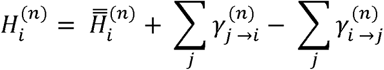

In the viability check phase the herd size of the current and the previous year is checked for all agents. If an agent’s herd fails to satisfy the threshold criteria for two consecutive years (i.e.,*H_i_^(n)^* < *θ* and *H_i_*^(*n*-1)^ < 0), then the agent and its herd are removed from the simulation and become unavailable for interaction and transfers.

This modeling assumption is based upon the fact that, among pastoralist societies, individuals whose herds are below a sustainability threshold typically keep their remaining cattle but are “removed” from the social system; they leave the group and find alternative ways (e.g., hunting, gathering, bee-keeping) to survive outside the network of herders. Surviving agents’ herds will then restart a typical yearly cycle with random growth phase until the simulation reaches the end.

**ODD protocol.**

### Model Description

The model description offered below follows the standardized ODD protocol for describing individual and agent based models (Grimm et al. 2006) and is based on Aktipis et al (2011). The model is a multiplayer-social network extension of the original model published in Aktipis et al (2011).

### Purpose

Here we use an agent-based model of wealth transfers within ecologically realistic conditions to investigate the viability of two sets of cooperative rules: one characterized by account keeping and the other characterized by risk pooling norms of need-based transfers. We then investigate how these two rules affect overall resource stock survivorship and the variability of survivorship within socially structured populations (i.e., agents interactions are constrained by the topology of social network they are part of).

### State variables and scales

In this model time is represented as discrete. Space is not explicitly modeled. Resource stock growth dynamics and volatility are implemented with global variables while the resource stock size and giving/asking rules are agent variables (Table 1 - below). During each time period, agents execute the commands described in the schedule.

**Table 1.** State variables and scales.

| Entity | State variable | Description |
| --- | --- | --- |
| Global | Growth rate | Amount by which resource stocks grow each year |
|  | Volatility rate | Likelihood of a negative event (e.g., drought) |
|  | Volatility size | Decrease in resource stock size resulting from negative event |
|  | Min. resource stock size | The minimum viable resource stock size |
|  | Max. resource stock size | The maximum resource stock that can be maintained |
|  | Correlation of shocks | Probability of correlated shocks |
| Agents | Resource stock size | Number of resources in agent's resource stock |
| | Net received list | Number of resources received minus given to $i$ partner (i.e., a list with length = $N$) |
|  | Asking threshold | The threshold below which agents ask for resources |
|  | Generosity | The likelihood of giving if asked and able (need-based) or giving to partner in good standing (account-keeping) |
| Need-based only | Giving threshold | The threshold below which agents will no longer give resources |
| Debt-based only | Credit size | The amount of credit granted to partner |
|  | Tolerated delay | The number of periods an agent will tolerate not being repaid by a partner before placing that partner in bad standing |
| | P good standing list | If $i$ partner is in good standing, means that they have not exceeded tolerated delay in past transfers (i.e., a list with length = $N$) |
|  | Repayment probability | The likelihood of repaying each partner during each time period |

### Process overview and scheduling

This model proceeds in discrete time steps, and entities execute procedures according to the following ordering:

1. For each actor, resource stocks change in size:
  a. Resource stocks increase in size according to growth rate
  b. Resource stocks decrease in size by volatility size (as a percent of total holdings) according to volatility rate
  c. If resource stock size is above resource stock max it is set to resource stock max
  d. Resource stock size is rounded to nearest integer
2. Couples of agents are selected from the social network following one of two possible rules:
  a. randomly select an agent as first player, then randomly select another agent among those which are linked to the first one
  b. randomly select first agent, then select the wealthiest agent among those linked to the first (i.e., following a “selective” asking process instead of a random one)
3. Requests are made: If giving is need-based, requests are made if resource stock size is below resource stock min If giving is account-keeping-based, requests are made if resource stock size is below resource stock min
4. Transfers are made: If giving is need-based, requests are fulfilled to the extent possible keeping the resource stock size of the giver above resource stock min If giving is account-keeping-based: If resources have been previously transferred from the partner to the actor, the actor transfers net received resources to their partner according to repayment prob. If a new request was made, actors give if two conditions are met: The debt has not existed for longer than tolerated delay The amount transferred cannot exceed the credit size extended to the partner. All actors update net received to reflect transfers
5. Actors removed from the population if two consecutive rounds occur where resources holdings are below resource stock min.
6. Age of actors incremented by 1

### Design concepts

Emergence:

In this model, risk pooling emerges from interactions between agents.

Prediction:

Agents in this model lack the ability to predict outcomes of future environmental variability or future social interactions. They do not integrate information across time periods.

Sensing:

Agents receive requests from their interaction partners and are able to examine their own resource holdings before fulfilling requests.

Interaction:

Agents interact by making and fulfilling requests for resources.

Stochasticity:

Resource stock growth and environmental volatility both have stochastic components.

Observation:

Reported data are averaged from 10,000 runs. Simulations were run until all agents were removed from the population (i.e., dropped below the viability threshold for more than 2 consecutive time periods).

### Initialization

All runs were initialized according to default parameters in the table 2 below.

**Table 2.** Initialization.

| Entity | State variable | Initial/Default value | Units |
| --- | --- | --- | --- |
| Global | Growth rate | 3.4 (SD: 2.53) | % current resource stock |
|  | Volatility rate | 10 | % per year |
|  | Volatility size | 30 (SD: 10) | % current resource stock |
|  | Min. resource stock size | 64 | Number resources |
|  | Max. resource stock size | 600 | Number resources |
|  | P of correlated shocks | {0, 0.1, 0.2, 0.3, 0.4, 0.5, 0.6, 0.7, 0.8, 0.9, 1} | probability |
| Agents | Resource stock size | 70 | Number resources |
|  | Net received list | [0, ..., 0] with length = N | Number resources |
|  | Asking threshold | 64 | Number resources |
|  | Generosity | 100 | % likelihood |
| Need-based only | Giving threshold | 64 | Number resources |
| Debt-based only | Credit size | 10 | Number resources |
|  | Tolerated delay | 5 | Years |
|  | P good standing list | [1, ..., 1] with length = N | Boolean value [0,1] |
|  | Repayment probability | 100 | % likelihood |

### Input

In order to make our model of the Maasai pastoral system as realistic as possible, the following parameter values and assumptions about resource dynamics were based on existing scholarship (Dahl & Hjort, 1976).

### Growth rate

We used a 3.4% growth rate with an SD of 2.53 based on Dahl and Hjort’s (1976:66) estimate the growth rate in “normal” conditions to be 3.4%, with a maximum possible growth rate of roughly 11% and a minimum of approximately -6% (in the diminishing resource stocks example). Dahl and Hjort estimates are based on both empirical evidence and analytical modeling.

### Resource stock size

Initial resource stock sizes in our model were 70, with a minimum of 64 and a maximum of 600. These values were derived from Dahl and Hjort (1976:178) who state that a resource stock of 64 resources is sufficient to sustain a reference family. Resource stock sizes described in the text range from 60-100 cows and resource stocks larger than 600 are not considered viable (Dahl and Hjort, 1976:158).

### Volatility

We used a volatility rate of 0.1, meaning that on average a disaster (e.g., drought or disease) occurred every 10 years. In our model, this disaster reduced the resources resource stock by 30% on average, with a SD of 10%. Dahl and Hjort (1976:114-130) note that these disasters occur approximately every 10-12 years based on empirical data, and that the population decline (during disasters that occur every 10 years) should not be more than approximately 28%, based on analytical models.

## References

1 Aktipis, A. (2016), Principles of cooperation across systems: from human sharing to multicellularity and cancer. Evol Appl, 9: 17–36. doi:10.1111/eva.12303

2 Cronk, Lee, Colette Berbesque, Thomas Conte, Matthew Gervais, Padmini Iyer, Brighid McCarthy, Dennis Sonkoi, Cathryn Townsend, and Athena Aktipis. Managing risk through cooperation: Need-based transfers and risk pooling among the societies of the Human Generosity Project. *In press*. In Global Perspectives on Long-Term Community Resource Management, Ludomir R. Lozny and Thomas H. McGovern, eds. Springer.

3 Winterhalder, B. 2007. Risk and decision-making. In The Oxford Handbook of Evolutionary Psychology, edited by R. I. M. Dunbar and Louise Barrett, pp. 433–445. Oxford: Oxford University Press.

4 Dorfman, M. S. (2007). Introduction to Risk Management and Insurance, Prentice Hall.

5 Hruschka, D. J. Friendship: Development, Ecology, and Evolution of a Relationship (University of California Press, 2010).

6 Boyd, R. & Richerson, P. J. Why culture is common but cultural evolution is rare. Proc. Br. Acad. 88, 73–93 (1996).

7 Onnela, J. P. et al. Structure and tie strengths in mobile communication networks. Proc. Natl Acad. Sci. USA 104, 7332–7336 (2007).

8 Barabàsi, A. L. & Albert, R. Emergence of scaling in random networks. Science 286, 509–512 (1999).

9 Nowak, M. A., Tarnita, C. & Wilson, E. O. The evolution of eusociality. Nature 466, 1057–1062 (2010).

10 Boyd, R. & Richerson, P. J. The evolution of reciprocity in sizable groups. J. Theor. Biol. 132, 337–356 (1988).

11 Eshel, I. & Cavalli-Sforza, L. L. Assortment of encounters and evolution of cooperativeness. Proc. Natl Acad. Sci. USA 79, 1331–1335 (1982).

12 Bowles, S. Group competition, reproductive levelling, and the evolution of human altruism. Science 314, 1569–1572 (2006).

13 Ohtsuki, H., Hauert, C., Lieberman, E. & Nowak, M. A. A simple rule for the evolution of cooperation on graphs and social networks. Nature 441, 502–505 (2006).

14 Apicella, C. L., Marlowe, F. W., Fowler, J. H., & Christakis, N. A. (2012). Social networks and cooperation in hunter-gatherers. Nature, 481(7382), 497–501.

15 Marlowe, F. W. The Hadza: Hunter-gatherers of Tanzania (University of California Press, 2010).

16 Nowak, M. A. Five rules for the evolution of cooperation. Science 314, 1560–1563 (2006).

17 Ash, J., & Newth, D. (2007). Optimizing complex networks for resilience against cascading failure. Physica A: Statistical Mechanics and its Applications, 380, 673–683.

18 Cronk, L. (2007). The Influence of Cultural Framing on Play in the Trust Game: A Maasai Example. Evolution and Human Behavior 28(5): 352–358.

19 Aktipis, C. A., Cronk, L., and de Aguiar, R. (2011). Risk-Pooling and Herd Survival: An Agent-Based Model of a Maasai Gift-Giving System. Human Ecology 39(2): 131140.

20 Dahl, G., and Hjort, A. (1976). Having Herds: Pastoral Herd Growth and Household Economy. Dept. of Social Anthropology, University of Stockholm, Stockholm.

21 Homewood, K. (2008). Ecology of African Pastoralist Societies. Oxford.

22 Hao, Y., et al., Need-based transfers on a network: a model of risk-pooling in ecologically volatile environments, Evolution and Human Behavior (2015), http://dx.doi.org/10.1016/j.evolhumbehav.2014.12.003

23 Mol, F. (1996). Maasai Language and Culture Dictionary. Maasai Centre, Lemek.

24 Perlov, D. C. (1987). Trading for influence: The social and cultural economics of livestock marketing among the highland Samburu of Northern Kenya. Ph.D. dissertation, anthropology, University of California - Los Angeles.

25 Aktipis, A., de Aguiar, R., Flaherty, A., Iyer, P., Sonkoi, D., & Cronk, L. (2016). Cooperation in an uncertain world: For the maasai of east africa, need-based transfers outperform account-keeping in volatile environments. Human Ecology, 44(3), 353–364. doi:http://dx.doi.org/10.1007/s10745-016-9823-z

26 Almagor, U. (1978). Pastoral Partners: Affinity and Bond Partnership Among the Dassanetch of South-West Ethiopia. Manchester Univ. Press, Manchester.

27 Bollig, M. (1998). Moral economy and self-interest: Kinship, friendship, and exchange among the Pokot (NW Kenya). In Schweizer, T., and White, D. (eds.), Kinship, Networks, and Exchange. Cambridge University Press, Cambridge. Cambridge University Press, Cambridge, pp. 137–157.

28 Bollig, M. (2010). Risk Management in a Hazardous Environment: A Comparative Study of Two Pastoral Societies (Vol. 2). Springer, New York.

29 Dyson-Hudson, N. (1966). Karimojong Politics. Clarendon, Oxford.

30 Flannery, K., Marcus, J., and Reynolds, R. (1989). The Flocks of the Wamani: A Study of Llama herders on the Punas of Ayacucho, Peru. Academic, New York.

31 Gulliver, P. (1955). The Family Herds: A Study of Two Pastoral Tribes in East Africa: the Jie and Turkana. International Library of Sociology and Social Reconstruction.

32 Iyer, Padmini. 2016. Risk Management Strategies of Male and Female Pastoralists in Karamoja, Uganda. Ph.D. dissertation, anthropology, Rutgers University.

33 Bramoullé, Y. and Kranton, R. Risk-sharing networks. Journal of Economic Behavior & Organization, 64, 3–4, 2007. https://doi.org/10.1016/jjebo.2006.10.004.

34 Pereda, M., Zurro, D., Santos, J. I., Briz i Godino, I., Álvarez, M., Caro, J., & Galán, J. M. (2017). Emergence and Evolution of Cooperation Under Resource Pressure. Scientific Reports, 7, 45574. http://doi.org/10.1038/srep45574

35 Wilensky, U. (1999). NetLogo. http://ccl.northwestern.edu/netlogo/. Center for Connected Learning and Computer-Based Modeling, Northwestern University, Evanston, IL.

36 Cronk, Lee. 2015. “Neighboring”: a preliminary look at generosity and mutual aid among ranchers in the American Southwest (http://humangenerosity.org).

37 Steinbacher, M., Steinbacher, M. & Steinbacher, M., Robustness of banking networks to idiosyncratic and systemic shocks: a network-based approach. (2016) Journal of Economic Interaction and Coordination, 11: 95. doi:10.1007/s11403-014-0143-3

38 Aktipis, C. A. (2004). Know When to Walk Away: Contingent Movement and the Evolution of Cooperation in Groups. Journal of Theoretical Biology 231(2): 249–260.

39 Aktipis, C. A. (2004). Know When to Walk Away: Contingent Movement and the Evolution of Cooperation in Groups. Journal of Theoretical Biology 231(2): 249–260.

40 Barclay, P. (2013). Strategies for Cooperation in Biological Markets, Especially for Humans. Evolution and Human Behavior 34(3): 164–175.

41 Barclay, P., and Willer, R. (2007). Partner Choice Creates Competitive Altruism in Humans. Proceedings of the Royal Society B: Biological Sciences 274(1610): 749753.

42 Campennì, M., & Schino, G. (2014). Partner choice promotes cooperation: the two faces of testing with agent-based models. Journal of theoretical biology, 344, 49–55.

43 Noe, R., and Hammerstein, P. (1994). Biological Markets: Supply and Demand Determine the Effect of Partner Choice in Cooperation, Mutualism and Mating. Behavioral Ecology and Sociobiology 35: 1–11.

44 Bullmore, E., & Sporns, O. (2012). The economy of brain network organization. Nature Reviews Neuroscience, 13(5), 336–349.

45 Eriksen, K. A., Simonsen, I., Maslov, S., & Sneppen, K. (2003). Modularity and extreme edges of the Internet. Physical review letters, 90(14), 148701.

46 Guimera, R., Mossa, S., Turtschi, A., & Amaral, L.N. (2005). The worldwide air transportation network: Anomalous centrality, community structure, and cities’ global roles. Proceedings of the National Academy of Sciences, 102(22), 7794–7799.

47 Garas, A., Argyrakis, P., & Havlin, S. (2008). The structural role of weak and strong links in a financial market network. The European Physical Journal B-Condensed Matter and Complex Systems, 63(2), 265–271.

48 Bettencourt, L. M., Lobo, J., Helbing, D., Kühnert, C., & West, G. B. (2007). Growth, innovation, scaling, and the pace of life in cities. Proceedings of the National Academy of Sciences, 104(17), 7301–7306.

49 Ripley, A. The unthinkable: Who survives when disaster strikes - and why. Harmony, 2009.

50 Solnit, R. A paradise built in hell: The extraordinary communities that arise in disaster. Penguin, 2010.

## Appendix References

● Aktipis, A. (2016), Principles of cooperation across systems: from human sharing to multicellularity and cancer. Evol Appl, 9: 17–36. doi:10.1111/eva.12303

● Aktipis, C. A., Cronk, L., and de Aguiar, R. (2011). Risk-Pooling and Herd Survival: An Agent-Based Model of a Maasai Gift-Giving System. Human Ecology 39(2): 131–140.

● Hao, Y., et al., Need-based transfers on a network: a model of risk-pooling in ecologically volatile environments, Evolution and Human Behavior (2015), http://dx.doi.org/10.1016/j.evolhumbehav.2014.12.003

● Wilensky, U. (2005). NetLogo Preferential Attachment model. http://ccl.northwestern.edu/netlogo/models/PreferentialAttachment. Center for Connected Learning and Computer-Based Modeling, Northwestern University, Evanston, IL.

● Albert, Réka; Barabási, Albert-László (2002). Statistical mechanics of complex networks. Reviews of Modern Physics. 74 (1): 47–97.

● Dahl, G., and Hjort, A. (1976). Having Herds: Pastoral Herd Growth and Household Economy. Dept. of Social Anthropology, University of Stockholm, Stockholm.

● Grimm, V., Berger, U., Bastiansen, F., Eliassen, S., Ginot, V., Giske, J., Goss-Custard, J., Grand, T., Heinz, S., and Huse, G. (2006). A Standard Protocol For Describing Individual-Based and Agent-Based Models. Ecological Modeling 198: 115–126.

